# The role of motion in the neural representation of social interactions in the posterior temporal cortex

**DOI:** 10.1101/2022.08.01.502134

**Authors:** Julia Landsiedel, Katie Daughters, Paul E. Downing, Kami Koldewyn

## Abstract

Humans are an inherently social species, with multiple focal brain regions sensitive to various visual social cues such as faces, bodies, and biological motion. More recently, research has begun to investigate how the brain responds to more complex, naturalistic social scenes, identifying a region in the posterior superior temporal sulcus (SI-pSTS; i.e., social interaction pSTS), among others, as an important region for processing social interaction. This research, however, has presented images *or* videos, and thus the contribution of motion to social interaction perception in these brain regions is not yet understood. In the current study, 22 participants viewed videos, image sequences, scrambled image sequences and static images of either social interactions or non-social independent actions. Combining univariate and multivariate analyses, we confirm that bilateral SI-pSTS plays a central role in *dynamic* social interaction perception but is much less involved when ‘interactiveness’ is conveyed solely with static cues. Regions in the social brain, including SI-pSTS and extrastriate body area (EBA), showed sensitivity to both motion and interactive content. While SI-pSTS is somewhat more tuned to video interactions than is EBA, both bilateral SI-pSTS and EBA showed a greater response to social interactions compared to non-interactions and both regions responded more strongly to videos than static images. Indeed, both regions showed higher responses to interactions than independent actions in videos and intact sequences, but not in other conditions. Exploratory multivariate regression analyses suggest that selectivity for simple visual motion does not in itself drive interactive sensitivity in either SI-pSTS or EBA. Rather, selectivity for interactions expressed in point-light animations, and selectivity for static images of bodies, make positive and independent contributions to this effect across the LOTC region. Our results strongly suggest that EBA and SI-pSTS work together during dynamic interaction perception, at least when interactive information is conveyed primarily via *body* information. As such, our results are also in line with proposals of a third visual stream supporting dynamic social scene perception.

## 1. Introduction

Social interactions are centrally important in the human experience. As inherently social beings, humans observe and engage in numerous social interactions every day and base much of their understanding of others’ actions and motivations on the information they glean from these social encounters. Despite the importance of interactive information in building our understanding of others, prior neuroimaging work has largely focused on the visual perception of individuals and the social cues they convey rather than on multi-person interactions. For example, several discrete areas in the brain have been characterised that are selectively responsive to socially relevant cues about single people. These include the fusiform face area (FFA), occipital face area (OFA), and a face-selective region of the superior temporal sulcus (fSTS), that are selective for faces (Gauthier et al., 2000; Kanwisher & Yovel, 2006; Phillips et al., 1997; Puce et al., 1998); the extrastriate body area (EBA) and fusiform body area (FBA) that are selective for bodies (Downing & Peelen, 2011; Peelen & Downing, 2005a; Schwarzlose et al., 2005), a region in the superior temporal sulcus (STS) that is selective for biological motion (Grossman & Blake, 2002), and a region of the temporal parietal junction (TPJ) that is selective for mentalising (Saxe & Kanwisher, 2003). While those studies have identified candidate regions that likely play some role(s) in perception of social interactions, they do not speak directly to the question of brain regions involved in processing social interactions *per se.*

Indeed, recent work suggests that observed interactions are perceived and processed in a way that extends beyond the sum of perceiving multiple individuals (Abassi & Papeo, 2020, 2021; Bellot et al., 2021; Isik et al., 2017; Masson & Isik, 2021; Walbrin et al., 2018; Walbrin & Koldewyn, 2019). Studies focused on dynamic interactions (Isik et al., 2017; Masson & Isik, 2021; Walbrin et al., 2018) have highlighted a region of the posterior superior temporal sulcus (pSTS) as being selectively involved in the perception of observed social interactions. Notably, this region is engaged even by stimuli that contain minimal or even no realistic human visual features. For example, the pSTS is activated by interactions performed by animated shapes (Castelli et al., 2000; Walbrin et al., 2018), point-light animations (Centelles et al., 2011; Sapey-Triomphe et al., 2017; Walbrin et al., 2018), and animated mannikins (Georgescu et al., 2014). Noting that the lateral temporal cortex is a large region with a complex profile of responses to social stimuli (Allison et al., 2000; Deen et al., 2015), it is important that neuroimaging evidence shows that the interaction-selective pSTS region (SI-pSTS) is anatomically and functionally dissociated from nearby regions that are also implicated in social processes. Specifically, Isik et al (2017) demonstrated this by functionally localising, at individual level, the STS interaction-selective region (SI-pSTS) along with motion-selective hMT+, the face-selective (fSTS) region, and the mentalizing-selective region of TPJ. Anatomical overlap amongst these regions was minimal; and importantly, selectivity for observing simple social interactions was significantly greater in the SI-pSTS region.

However, the SI-pSTS is not the only posterior region implicated in visual interaction perception. Recent studies using static images of dyads who are either facing each other, or not, has suggested that extrastriate body area (EBA), rather than the pSTS, is more uniquely engaged by facing human dyads (Abassi & Papeo, 2020, 2021). In addition, two studies using dynamic dyads that gesture and/or move towards vs. away from each other find that the EBA, as well as pSTS, plays a role in processing social interactions (Bellot et al., 2021; Walbrin & Koldewyn, 2019).

In sum, these studies point to candidate brain regions that contribute to interaction perception, and at the same time they suggest that the distinction between moving and static depictions of interactions may be an important moderating variable. Previous studies of interaction perception have almost universally presented either static or dynamic stimuli but have not directly assessed the role of motion. Therefore, the present paper investigates the extent to which the interaction selectivity of these brain regions depends on the presence of dynamic information, as this is critical for building a full picture of how interactive information is processed and understood in the human brain.

There are good reasons to suspect that motion plays a key role in social interaction perception and that regions sensitive to interactive information should also be sensitive to (at least some) motion cues. First, as outlined by Pitcher and Ungerleider (2021), the STS as a whole contains body and face selective regions that are more responsive to moving than static faces and bodies. Second, although observers are exquisitely sensitive to static cues to interaction such as facing direction, proximity, and touch in static scenes (Zhou et al., 2019), interactions by their very nature unfold dynamically. Thus, it is intuitive to expect that at least some regions involved in processing social interactions will do so preferentially in response to dynamic stimuli. Indeed, Pitcher and Ungerleider (2021) have proposed a “third” visual pathway that is specialised for dynamic social perception, a pathway that stretches from V1, through motion-selective middle temporal cortex (hMT+) and runs down the length of the STS. According to that proposal, both EBA and SI-pSTS regions would be expected to show motion sensitivity, although SI-pSTS might be expected to show greater sensitivity to complex social content.

Accordingly, in the present study we factorially manipulated both “interactivity” and motion to identify the contribution of both factors, and the potential interactions between them, on brain responses (see Figure 1). Stimuli included social interactions as well as non-interactive scenes depicting independent actions that were matched on high-level perceptual features such as actor identity and gender, and scene and object contents, as well as on motion energy and physical distance. Orthogonally, in a design similar to that of Pitcher et al. (2011; see also Downing et al., 2006 and Hasson et al., 2008), scenes were presented as dynamic movies, or as still frames that were extracted from the interaction movies and that were either presented in the correct sequence, in a fixed random order, or else as a single “key” image.

**Figure 1.**
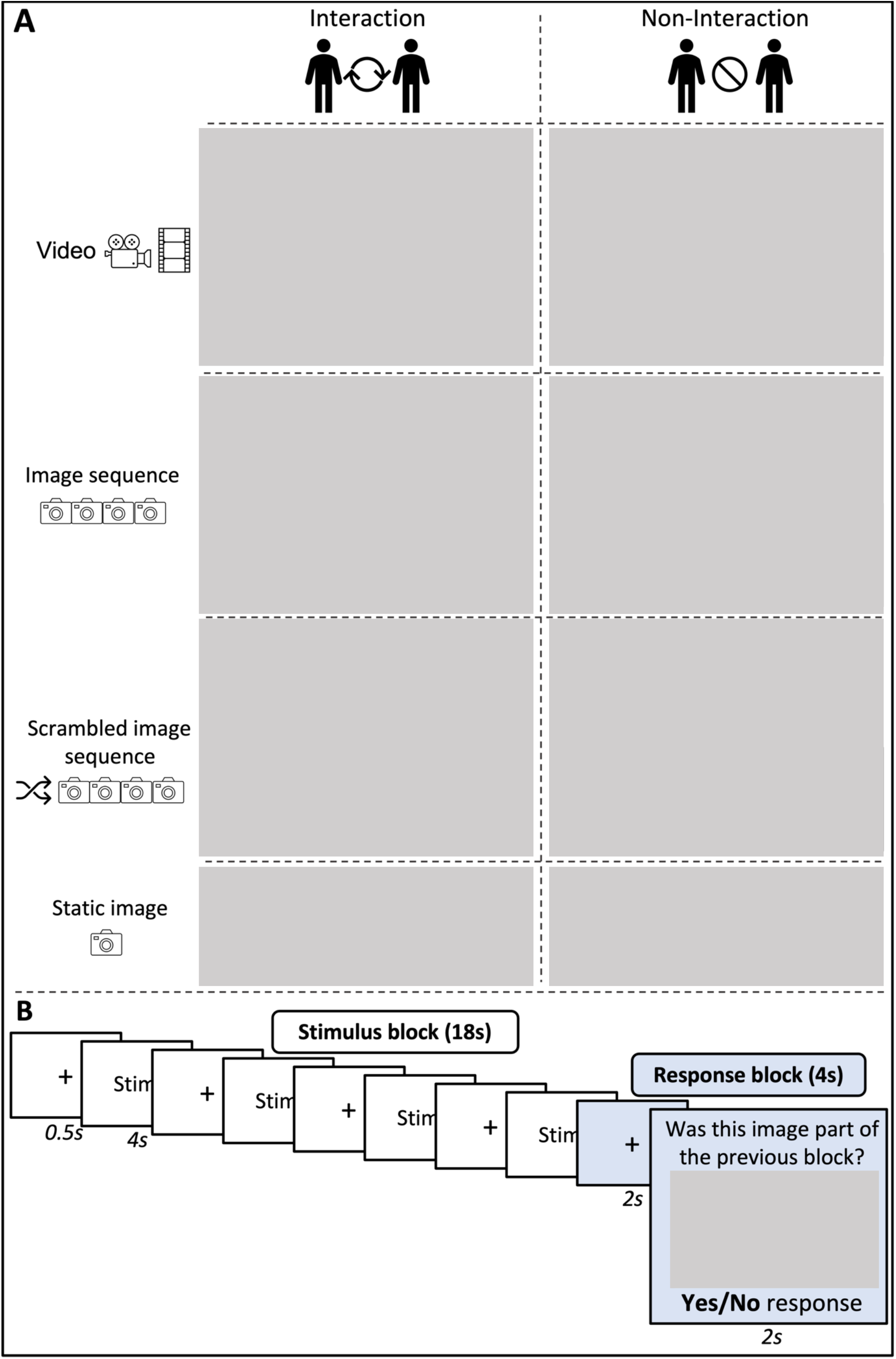
A: Illustration of 2 (Scene) x 4 (Motion) design; B: Illustration of the main task: an 18-second stimulus block is followed by a 4-second response block.

A further important advance of the present work over many previous studies is that we presented social interactions that were depicted in a range of naturalistic everyday settings and using a variety of interactive cues. A small number of previous studies have taken a similar naturalistic approach to related issues. One such study presented short video clips of computer animations depicting a range of social (e.g., faces, bodies and social interactions) and non-social (e.g., objects, houses and non-social interactions) features, finding that although the posterior temporal cortex in general responded more to social compared to non-social stimuli, the pSTS showed a significant increase in response for all eight identified social features, including social interaction (Lahnakoski et al., 2012). Taking this one step further, two recent studies have analysed data collected while participants viewed TV episodes or excerpts from movies (Masson & Isik, 2021; Wagner et al., 2016). Wagner et al. (2016) found that medial prefrontal cortex (mPFC) was most involved in processing naturalistic social interactions and speculated that this was because social interactions trigger spontaneous mentalising processes. Masson and Isik (2021), however, found bilateral pSTS to be most involved in social interaction perception. They assessed the contribution of both mentalising processes and social interaction perception, concluding that social interactions explained unique variance in the response of the pSTS to naturalistic scenes in a way that was not replicated in “canonical” ToM regions, including mPFC. While some of this work has assessed motion as one perceptual feature among many, it leaves open key questions about the ways that motion drives the response to interactions in social perception regions.

The existing literature outlined above points to a diverse set of known temporal and occipito-temporal brain regions that may contribute to social interaction perception. Accordingly, we adopted a functional localiser approach (e.g., Kanwisher, 2017; Saxe et al., 2006), identifying key regions of interest with independent localiser datasets in each participant, and measuring the response of these regions to the interactive and control stimuli described above (see Figure *2*). These regions of interest (ROIs) included: SI-pSTS, defined by greater response to interacting than non-interacting point-light dyads (Isik et al, 2017; Walbrin et al, 2018); EBA and its ventral counterpart the fusiform body area (FBA; Peelen and Downing, 2005a; Schwarzlose et al., 2005) localised by a selective response to human bodies; a TPJ ROI implicated in mentalising, localised by contrasting selected epochs within a brief movie (Jacoby et al., 2016); and motion-selective hMT+, localised with a contrast of simple visual motion vs a static control (Tootell et al., 1995).

**Figure 2.**
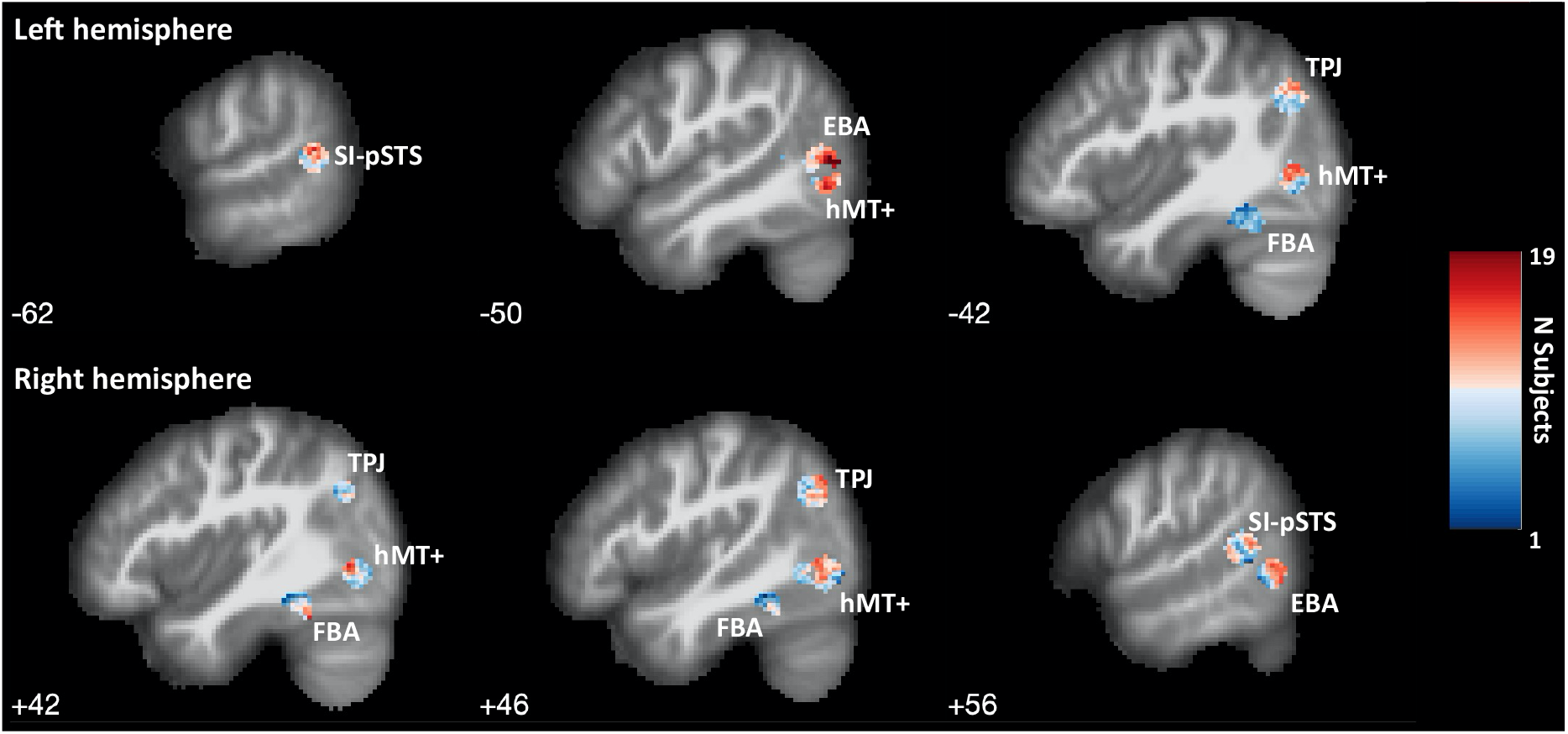
Functionally-defined regions of interest for univariate analyses of responses to social interactions displayed through a sagittal view heatmap of subject-specific ROI overlap (x-coordinates in MNI space).

For two of these regions, clear hypotheses about the individual and combined effects of motion and interaction content are possible. First, we hypothesised that the SI-pSTS ROI would be more responsive to interactive than non-interactive scenes and would also be more responsive to dynamic than static conditions. However, we also expected that these two factors would interact such that the difference between social conditions would be largest in the video condition; that is, that the SI-pSTS would show higher interaction sensitivity when scenes are presented dynamically. We also expected that the SI-pSTS would be more responsive to interactive information when static scenes in sequences were shown in a coherent order, with intact implied motion cues, than when presented in a scrambled order. Second, and in contrast to SI-pSTS, we hypothesised that EBA would show a main effect of motion (see Downing et al., 2001), but only weak sensitivity to interactive information. We did not expect EBA to be particularly sensitive to interactive information in dynamic scenes, but instead expected that EBA might be more responsive to interactive cues conveyed through static body cues, in line with prior work (Abassi & Papeo, 2020, 2021).

Further occipito-temporal regions of interest served as controls to determine the spatial and functional specificity of responses to social interactions. The bilateral TPJ ROI identified in a theory-of-mind contrast was included because it is spatially very near SI-pSTS, and is clearly implicated in social cognition, but (based on previous findings) was not expected to be especially sensitive to motion nor to the combined effects of motion and social interaction (Walbrin et al., 2019; Walbrin et al., 2020). Given the key role of human bodies in depicting natural interactions, FBA responses were compared to those of EBA. Previous dissociations in the functional properties of these two body-selective regions (Ewbank et al., 2011; Schwarzlose et al., 2008; Taylor & Downing, 2011) and their location in distinct neuroanatomical pathways (Pitcher and Ungerleider, 2021) indicated the possibility of distinct responses to social interactions between these counterpart regions. Finally, as a reality check, we included hMT+ as a region that is generally responsive to motion and optic flow, which we expected should be clearly modulated by our motion manipulation without showing sensitivity to the interactiveness of the scenes.

We used these functional localisers to test our hypotheses with two complementary analytical approaches. First, standard univariate analyses revealed the magnitude of response to the conditions of the main experiment in each ROI. These analyses indexed the degree to which individually defined social brain regions are differentially activated, on the whole, to social interactions. Second, we adopted a multivoxel regression approach to assess how, over a broad occipitotemporal region, the various specific selectivities that were measured in the localiser data (e.g., to static bodies, or to simple visual motion) jointly predicted the response to social interactions. These analyses allowed a more continuous view of selectivity, capitalising on local variation in response profiles rather than relying on binary classifications of regions as selective or not for a given property.

## 2. Methods

### 2.1 Participants

Twenty-four right-handed participants were recruited to take part in this study, two of which were removed from the analyses due to excessive head motion (see 2.6 *MRI pre-processing* section for details). Thus, the final sample consisted of 22 participants (mean age = 21.59, SD = 2.11, 6 male). All participants had normal or corrected to normal vision, provided written informed consent, and were fully debriefed at the end of the study. Participants received monetary compensation for taking part. The study was approved by the Ethics Committee of the School of Psychology at Bangor University and was pre-registered on AsPredicted ID 52482 on 18/11/2020.

### 2.2 Design & Procedure

To investigate sensitivity to different levels of motion in social interactions and noninteractions, the main experimental task consisted of a 2 (Scene: Social interaction vs non-interaction) × 4 (Motion: Dynamic videos, image sequences, scrambled image sequences, static images) repeated measures fMRI block design (see Figure 1A).

Experimental conditions were presented in blocks of stimuli followed by a short response block. Stimulus blocks contained four 4-second stimulus trials, each preceded by a 0.5-second black fixation cross on a grey background (18 seconds total). For video stimuli, four 4-second videos were shown. For image sequences, four sequences of four 1-second images each were shown in the correct order. For scrambled image sequences, four sequences of the same quadruplet 1-second images from the image sequence blocks were shown in a consistent pseudorandom order. For static images, four images were shown for 4-seconds each.

Every stimulus block was followed by a 4-second response block: 2 seconds fixation before a unique response image probe was shown for 2 seconds (thus 22-second total experimental block duration). Participants were asked to identify, within the 2-second presentation time, whether the scenario depicted in the response image had been part of the stimulus presentation phase of the current block with a dichotomous yes/no response. This was coded such that there was a 50/50 chance of the scenario having been shown. For an illustration of this procedure, refer to Figure 1B.

Functional data were acquired across four runs lasting 400 seconds each. Each functional run contained two blocks of each experimental condition (22 seconds each, 16 blocks per run) and three rest blocks (16 seconds each) at the beginning, middle, and end of the run. The order of blocks in each run was palindromic and pseudorandomised across runs such that each of the eight blocks was presented at a different point in each run. A total of eight runs were created, four of which were chosen at random for each participant, but all runs were shown an equal number of times across the study.

The experiment was created using PsychoPy 3 (Peirce et al., 2019) running on Linux Ubuntu 18.04.

### 2.3 Stimuli

Novel stimuli were created for this study and validated in an independent study (see Appendix 1). Sixty 4-second video clips were created of various everyday scenarios. Each scenario depicted two individuals either engaging in a social interaction or acting independently (non-interactions). In total, there were 30 interactive and 30 non-interactive scenarios. The social interactions and non-interactions were matched as closely as possible, such that they both contained the same action, props, and (so far as possible) maintained the same average physical distance between the two individuals. Scenarios were acted out by four different actor pairings (two female/female pairs, one male/male pair and one female/male pair) captured in eight different geographical locations. Following the approach by Grezes et al. (2007), social interaction and non-interaction videos were analysed for differences in motion (see Appendix 2). A paired sample t-test confirmed that the two conditions did not differ in their overall motion energy, *t*(29) = −1.61, *p* = 0.12. Descriptively, non-interaction videos contained slightly more movement than social interaction videos.

All other stimulus conditions (image sequences, scrambled image sequences, static images) as well as response image probes were created from the video stimuli by selecting individual video frames. For image sequences, one frame was chosen to depict the beginning, two for the middle and one for the end of the scenario (approximately, at 0.5s, 1.5s, 2.5s, 3.5s). Specifically, images were taken from videos such that two individuals were identifiable in every image and for interactive scenarios so that the main interaction would be captured in the middle two frames. For scrambled image sequence blocks, the same quadruplet images from the image sequence stimuli were used but the order of the four images within a scenario were presented in a consistent pseudo-random order. Pseudo-randomisation was used to ensure that the scrambled sequence looked out of sync for each specific scenario. For static images, a specific key frame was identified for each video stimulus being the most representative for each social interaction or non-interaction scenario respectively. Finally, response image probes were selected to capture a moment that was suitably different than other image frames already taken from each scenario.

### 2.4 Localizer Tasks

For region-of-interest (ROI) analyses, all participants completed four localizer tasks in addition to the main experimental task. Three participants had already completed two or three of the four localizers during a previous study and did not complete these scans again. All localizer tasks were presented in Psychtoolbox 3.0.16 (Brainard, 1997; Kleiner et al., 2007) using MATLAB 2020.b (The MathWorks Inc.) running on a Linux Ubuntu 18.04 distribution stimulus computer.

A social interaction region localizer (Isik et al., 2017; Walbrin et al., 2018) was used to localize subregions of bilateral pSTS that are engaged by social interactions (SI-pSTS). Participants viewed 16-second blocks with videos of two point-light figures that are either interacting, not interacting/acting independently, or scrambled. Participants were instructed to simply watch the videos. Participants completed three runs of 2.5 minutes each; each run consisted of three 16-second rest blocks and two blocks per interaction type, one presented in either half of each run, in counterbalanced order with the other conditions. Interaction selective SI-pSTS ROIs were localised with the interaction > noninteraction contrast.

A second localizer (cf. Peelen & Downing, 2005a) was used to localize bilateral EBA and FBA. Participants completed two runs of a 1-back task. There were five 16-second rest blocks spread evenly across each run. Each of the four stimulus blocks (faces, bodies, chairs, and scenes) was presented once between each pair of rest blocks. Each stimulus block contained 24 stimulus exemplars drawn from a pool of 40 images. Stimuli were presented sequentially and appeared for 300 msec followed by a 700-msec ISI. Repetitions occurred twice per block. Both EBA and FBA were localized using the bodies > objects contrast.

A third localizer (Jacoby et al., 2016) was used to localize the TPJ. Participants were asked to attentively watch the Pixar short film ‘Partly Cloudy’ (2009, 5:49 minutes). The film scenes were coded by event type (mentalizing, pain, social, and control) and the contrast mentalizing vs. pain was used to localize bilateral TPJ. This approach has been demonstrated to robustly identify functional regions of interest involved in mentalising and other aspects of social cognition, that are in line with those identified with more standard blocked-design localizers.

Finally, we localized motion-sensitive human middle temporal cortex (hMT+; cf. Tootell et al., 1995). Participants completed one run of a passive viewing task in which they watched low contrast greyscale concentric circles either oscillating between expanding and contracting, or not moving. There were five 16-second rest blocks spread evenly across the run. Each of the two stimulus conditions (moving, static) was presented twice between each pair of rest blocks. The hMT+ was localised using the motion > non-motion contrast.

### 2.5 Behavioural Follow-up Task

To confirm that participants who took part in the study viewed the stimuli in a similar way to participants who completed the validation study described above, they were invited to complete a short behavioural task after their scan. During the task, participants were shown, one at a time, the key frame used in the static image condition and asked to rate how social they thought the image was, how visually interesting it was to look at and how positive they thought it was. Three paired t-tests were carried out to assess whether social interaction images were rated differently compared to non-interaction images.

### 2.6 MRI parameters and pre-processing

Images were acquired using a Philips Ingenia Elition X 3T scanner with a 32-channel head coil (Philips, Eindhoven, the Netherlands). For functional runs, a T2*weighted gradient-echo single-shot EPI pulse sequence (with SofTone mode noise reduction); TR = 2000ms, TE = 30ms, flip angle = 83°, FOV (mm) = 240×240×112, acquisition matrix = 80×78 (reconstruction matrix = 80); 36 contiguous axial slices in ascending order, reconstructed voxel size = 3×3×3mm^3^. Four dummy scans were discarded prior to image acquisition for each run.

For each participant, a high-resolution anatomical T1-weighted image acquired using a gradient echo, multi-shot turbo field echo pulse sequence, with a five-echo average; TR = 18ms, average TE = 9.8ms, in 3.2ms steps, total acquisition time = 338s, flip angle = 8°, FOV = 224×224, acquisition matrix = 224×220 (reconstruction matrix = 240); 175 contiguous slices, acquired voxel size (mm) = 1.0×1.0×2.0 (reconstructed voxel size = 1mm^3^).

Pre-processing steps included realignment and re-slicing, co-registration, segmentation, normalisation with 2mm isotropic voxel size in normalised MNI space, and spatial smoothing. Those steps, and general linear model (GLM) estimation were performed with SPM12 (fil.ion.ucl.ac.uk/spm/software/spm12). All SPM12 default pre-processing parameters were used except for the use of an initial 3mm FWHM Gaussian smoothing kernel. This smoothing kernel is recommended when intending to use ArtRepair toolbox (v5b, Mazaika et al., 2005). ArtRepair was used to detect and, when required, repair noisy volumes (volumes that contained 1.3% variation in global intensity or 0.5mm/TR scan-to-scan-motion). 21 subjects needed repairs in at least one run. Prior to first-level participant models, data was smoothed again using a 5mm FWHM kernel. This two-step procedure approximately results in the equivalent of data smoothed with a 6mm FWHM kernel.

Block durations and onsets for each experimental condition (per run) were modelled using a boxcar reference vector and convolved with a canonical hemodynamic response function (without time or dispersion derivatives) with a high-pass filter of 128 s and autoregressive AR(1) model. Repaired volumes were de-weighted in this analysis. First-level models for the functional localizers contained separate regressors for each condition. For the main experimental task, two separate first level models were used. Model 1 used eight separate regressors for each condition’s stimulus blocks. A further two regressors of no interest were used for response blocks which were collapsed across motion conditions for interaction and non-interaction blocks respectively. This model was used to extract percent signal change (PSC) for each experimental condition without including each condition’s response block in the PSC estimate. Model 2 used eight regressors for each condition respectively that included the combination of stimulus and response blocks for later time course visualisation. Across all tasks, rest blocks were modelled implicitly, and head motion was modelled using six nuisance regressors (three translation and three rotation)

### 2.7 Region of Interest (ROI) definition

ROIs were defined at the individual subject level using an established stepwise procedure (Julian et al., 2012, for a detailed description also see Walbrin et al., 2020). This approach uses group-level data to constrain the overall location of subject-specific ROIs; i.e., group-constrained ROI definition. Specifically, functional localizer group-level analyses were used to localize coordinates for the respective ROIs to define *initial* 8mm bounding spheres (see Appendix 3, Table A2). These were validated in a leave-one-subject-out (LOSO) iteration of the same group level analyses, resulting in a *subject-specific ROI search space* that included the overlap of the initial sphere and the respective LOSO group analysis. Due to the regional proximity of EBA and hMT+, any overlap of EBA and hMT+ search spaces was removed at this point. Finally, *subject-specific* ROIs were defined based on the respective first-level localizer contrasts masked by the *subject-specific ROI search space.* For all ROIs except FBA, the most activated 100 contiguous voxels (minimum threshold: T = 1) were included in the final ROI. As FBA is typically smaller in size than the other ROIs, is more variable, and can show significant overlap with FFA (Schwarzlose et al., 2005), we chose the 40 most activated voxels for FBA, rather than 100 (see Figure *2* for a heat map of subject-specific ROIs, visualised using bspmview toolbox; DOI: 10.5281/zenodo.168074, see also https://www.bobspunt.com/software/bspmview/)).

### 2.8 Percent signal change (PSC) analyses

For the main experimental task using Model 1, mean PSC per condition was estimated for each participant using Marsbar toolbox (Brett et al., 2002). Traditionally, we would extract PSC from unsmoothed data to due to our ROIs being functionally homogenous. However, the use of ArtRepair meant that PSC was extracted from minimally smoothed data (3mm FWHM). PSC values were extracted for all ten ROIs: bilateral SI-pSTS, TPJ, EBA, FBA and hMT+, for group analyses. Initially, for each hemisphere, an omnibus three-way 5 (Region: TPJ, SI-pSTS, EBA, hMT+, FBA) × 2 (Scene: Social interaction vs non-interaction) × 4 (Motion: Dynamic videos, image sequences, scrambled image sequences, static images) repeated-measures ANOVA was conducted. Subsequently, for each ROI, a 2 (Scene) × 4 (Motion) repeated measures ANOVA was run to analyse differences in PSC across all eight experimental conditions. Greenhouse-Geisser correction for sphericity violations were applied where required. A priori planned comparisons without Bonferroni correction included the difference between interactive vs non-interactive scenes for video and static motion conditions for bilateral SI-pSTS and EBA. Furthermore, post-hoc paired sample t-tests were conducted to follow-up any significant interaction effects. To account for multiple comparisons, the *p*-value was adjusted to *α* < 0.0125, taking into account four comparisons of interest, specifically the difference between interaction vs noninteraction for each motion condition. To examine interactions between scene and motion further, paired sample t-tests were also used to compare interaction ‘selectivity’ (calculated as the PSC difference between interactive and non-interactive scenes for each motion condition respectively) between motion conditions. For all paired t-test comparisons, effect sizes are expressed as Cohen’s d for repeated measures (*d_rm_*) which represents the mean difference standardized by the standard deviation of the difference scores corrected for the correlation between the measurements (Lakens, 2013).

To illustrate differences in the temporal dynamics between ROIs, the Marsbar toolbox (Brett et al., 2002) was also used to extract mean finite impulse response (FIR) time course data in PSC for each experimental condition. This analysis used GLM Model 2 to examine each conditions’ time courses including both stimulus and response blocks. The resulting time courses present the 22 s condition blocks as 11 time bins of 2 s; i.e. 1TR, each). As above, the plotted data represents interaction selectivity (PSC interactive minus non-interactive scenes) for each time bin and motion condition. Due to power constraints in the case of a 11 (Time bin) × 4 (Motion) repeated measures ANOVA, this data was *not* analysed statistically.

### 2.9 Exploratory multivariate analyses

Multivariate pattern analyses (MVPA) were used to explore the univariate results further, on the grounds that previous studies have identified greater sensitivity to condition differences in multivariate relative to univariate measures (Kamitani & Tong, 2005). An advantage of this approach is that it does not require a binary assignment of voxels as being (say) body-selective or not body-selective, but rather captures the voxelwise variation in selectivity over a broad region. Previously, this type of analysis has been used to disentangle the contribution of body- and motion-selectivity in the prediction of biological motion-selectivity in ventral regions, where it was demonstrated that biological motion selectivity was better predicted voxelwise by body selectivity than motion selectivity (Peelen et al., 2006).

Here, we adopted this approach to show how voxelwise variation in selectivity to bodies, to simple visual motion, and to point-light-interactions, might jointly explain the neural responses to interactions and non-interactions in the main experiment (see Figure 4 for an illustration of the steps involved). To this end, we examined a global lateral occipito-temporal cortex (LOTC) union combining SI-pSTS, EBA, and hMT+ (see Figure 3), which thus contained voxels that were highly category-selective for at least one of the three predictors. To create this global ROI, the same group level MNI coordinates used for individual EBA, hMT+ and SI-pSTS ROIs were used as centres of 12mm spheres. This radius was chosen to connect and capture all regions fully. For each hemisphere, these spheres were combined as the union across the three spheres.

**Figure 3.**
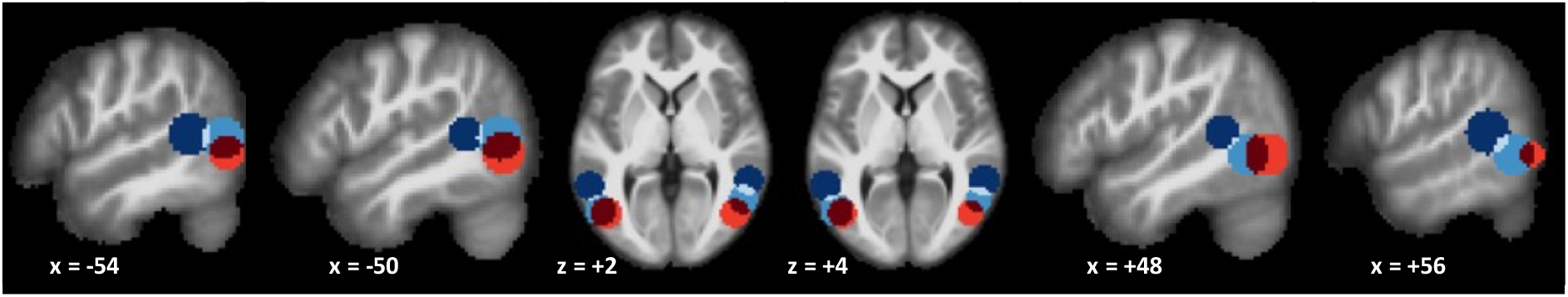
Axial and sagittal slices of bilateral combined 12mm spheres of SI-pSTS (dark blue), EBA (blue), hMT+ (light red) and their overlap (SI-pSTS-EBA overlap in light blue, EBA-hMT+ overlap in dark red)

**Figure 4.**
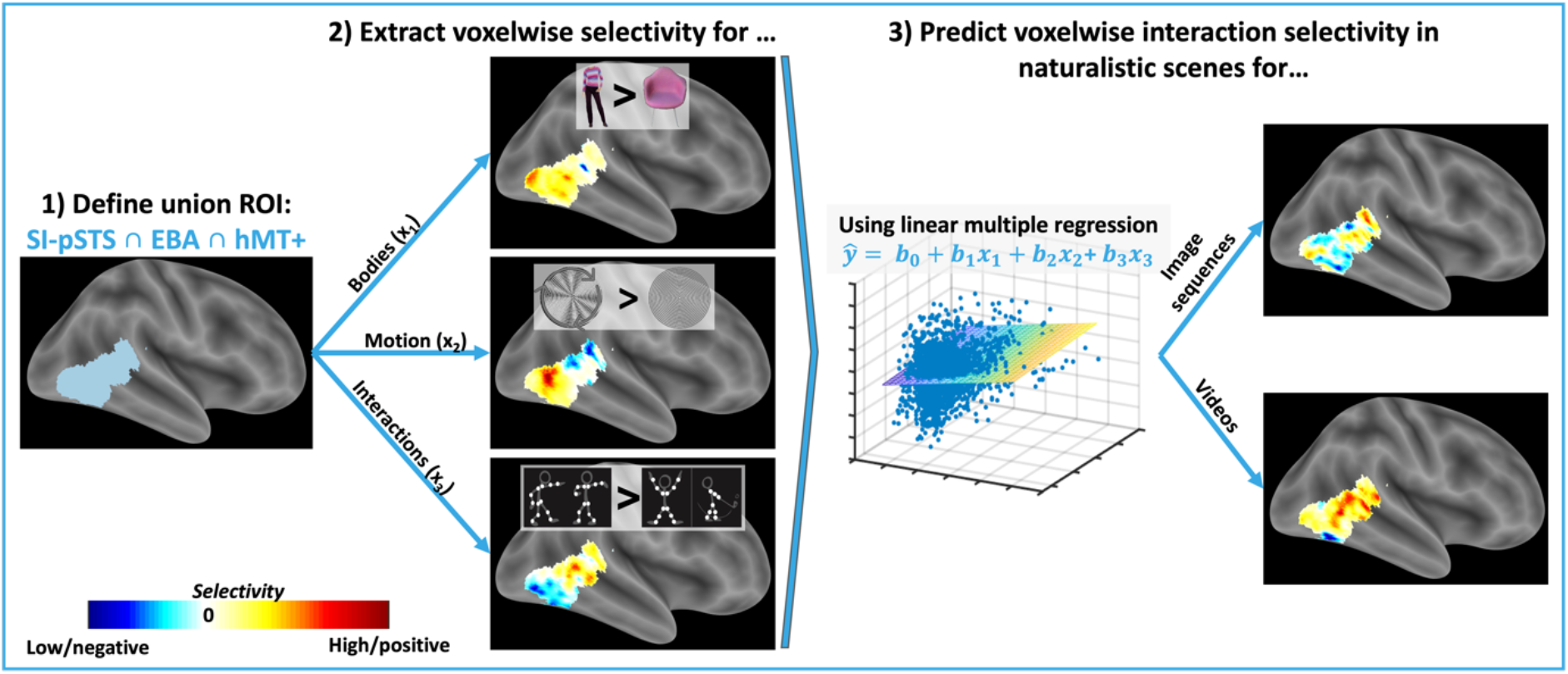
Illustration of exploratory MVPA analysis with linear multiple regression (for details see 2.9). The aim of the analysis was to measure the distributed contributions of three types of category selectivity to the perception of naturalistic social interactions in a broad lateral occipito-temporal (LOTC) region. This method does not require binary classification of each voxel as, for instance, either body or motion selective (or neither) but rather exploits continuous variation in these properties over the region. (1) For each hemisphere, a union ROI of SI-pSTS, EBA, and hMT+ was defined to capture voxelwise variation in selectivity for static bodies, simple visual motion, and point-light social interactions over the LOTC. (2) Voxelwise selectivity (difference in beta value estimates) for bodies, motion, and interactions was extracted for each participant and (3) entered as independent predictors into two separate linear multiple regressions to predict interaction selectivity in the main experimental task. One of these examined interaction selectivity (beta value difference interactive vs non-interactive scenes) in the Image Sequence condition, and the other in the Video condition. For illustration, the 3D scatterplot shows simplified exemplary group data using only two independent predictors (body and point-light interactions selectivity) in the regression model predicting interaction-selectivity for videos. Note: Selectivity maps are for illustrative purposes only and represent selectivity data averaged across the entire sample. Surface maps are voxel-to-surface projections created using bspmview; the analysis was conducted in voxel-space.

Using the localizer data, voxel-by-voxel patterns of body-, motion- and point-light-interaction-selectivity were extracted for each participant in this global LOTC ROI, bilaterally. These were calculated as the t-values of the difference for the respective contrast (e.g., bodies > chairs). Similarly, patterns of motion-dependent interaction selectivity (difference interaction > non-interactions for the image sequence and video conditions respectively) were also calculated and extracted. Subsequently, for each participant, this data was entered into two multiple regression analyses, one for predicting interaction selectivity in image sequences, and a similar one for full-cue videos. Each regression fitted a linear model including a constant term as well as body-, motion-, and point-light-interaction-selectivity as predictors. The resulting beta values for each predictor were aggregated across participants and compared against zero using one-sample t-tests. A predictor significantly greater from zero (e.g. for the bodies > chairs difference) could be interpreted to mean that that kind of selectivity (e.g. for bodies) made a unique contribution to variance in interaction-selectivity across the LOTC region.

## 3. Results

### 3.1 Behavioural

Replicating findings from the validation study, and in line with hypotheses, results demonstrated that 1) social interaction images were rated as more social compared to the non-interaction images (*t*(22) = 20.661, *p* < .001); 2) social interaction images were rated as more visually interesting compared to non-interaction images (*t*(22) = 11.170, *p* < .001); and 3) that social interaction images were rated as more positive compared to non-interactions images (*t*(22) = 13.377,*p* < .001).

### 3.2 Imaging

#### 3.2.1 Univariate ROI PSC analyses

##### Three-way ANOVAs

In the *right* hemisphere, the Region × Scene × Motion omnibus ANOVA found significant main effects of Region: *F(4,84)* = 12.72, *p* < .001, 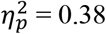, Scene: *F(1,21)* = 8.48, *p* < .01, 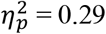, and Motion: *F(25.09,41.79)* = 122.37, *p* < .001, 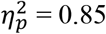. The Region × Scene: *F(2.80,58.88)* = 3.27, *p* = .03, 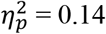, and Region × Motion: *F(3.28,68.91)* = 18.13, *p* < .001, 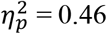, but not the Scene × Motion: *F(3,63)* = 2.11, *p* = .11, 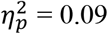, two-way interactions were significant as was the crucial three-way interaction: *F(6.37,133.73)* = 2.46, *p* = .03, 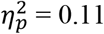.

In the *left* hemisphere, the Region × Scene × Motion omnibus ANOVA found significant main effects of Region: *F(4,84)* = 8.06, *p* < .001, 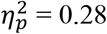, Scene: *F(1,21)* = 12.31, *p* < .01, 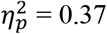, and Motion: *F(3,63)* = 107.55, *p* < .001, 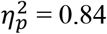. The Region × Motion: *F(3.32,69.69)* = 20.12, *p* < .001, 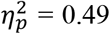, and the Scene × Motion: *F(3,63)* = 3.91, *p* = .01, 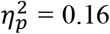, but not the Region × Scene: *F(4,84)* = 1.93, *p* = .11, 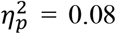, two-way interactions were significant. The crucial three-way interaction was marginally significant: *F(5.70,119.73,)* = 1.98, *p* = .08, 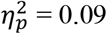.

Subsequently, these results were followed up for each ROI separately using Scene × Motion ANOVAs. Key results of these PSC analyses (see also Figure 5) are reported below (see Tables 1; for detailed descriptives please refer to Appendix 4, Tables A3 & A4). Additionally, time courses of motion-dependent interaction selectivity within our ROIs are presented to illustrate the temporal dynamics of the PSC response (see Figure 6).

**Figure 5.**
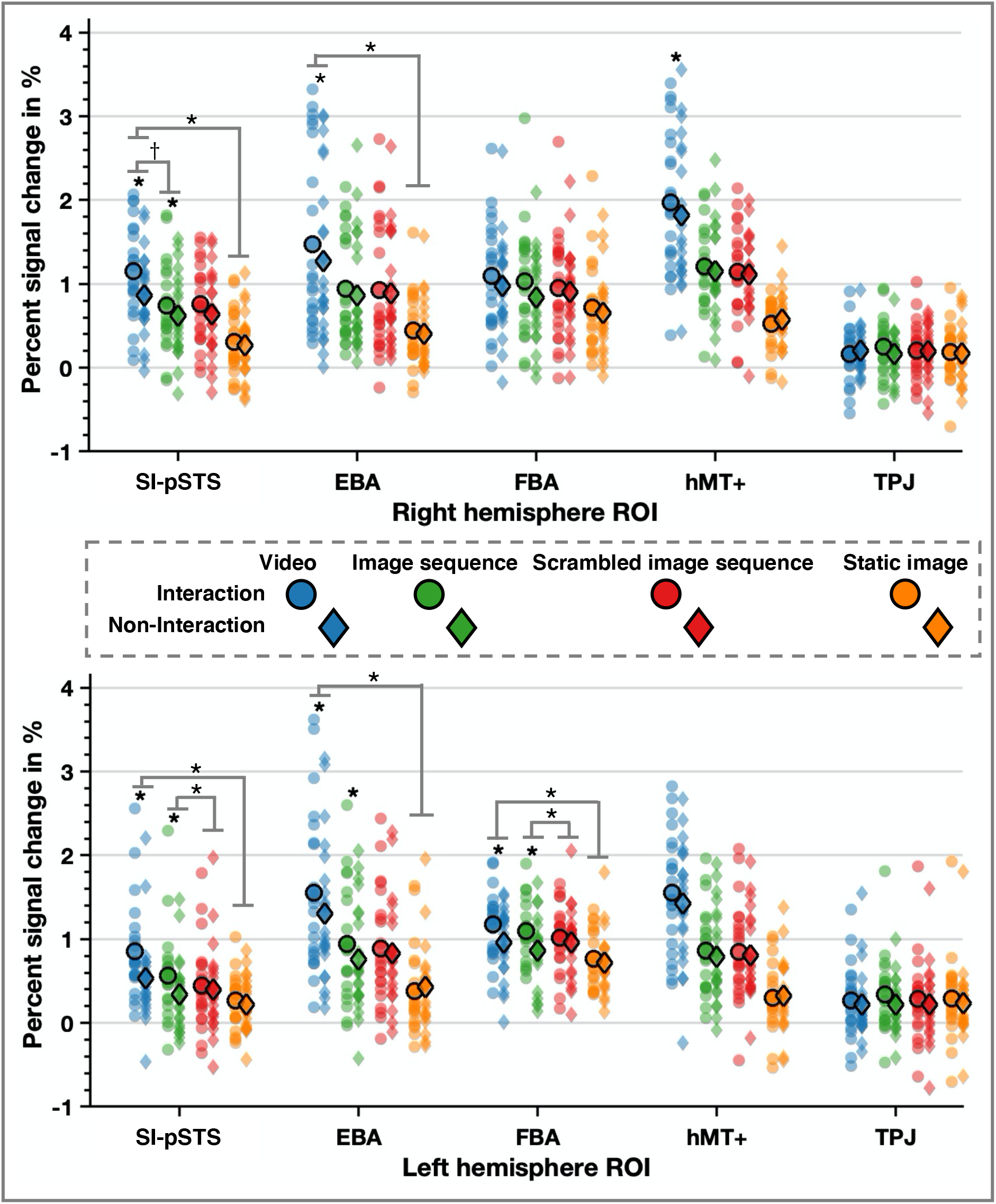
Percent signal change data displaying condition means (circle with bold edge) and data distribution for each ROI: top: right hemisphere, bottom: left hemisphere. Significant planned contrasts and Bonferroni corrected post-hoc paired t-test results between interactive and non-interaction conditions are marked using an asterisk. Significant differences between motion conditions in interaction selectivity are reflected by the grey lines and asterisk above the data distributions for the respective ROI (bilateral SI-pSTS, EBA, and left FBA only).

**Figure 6.**
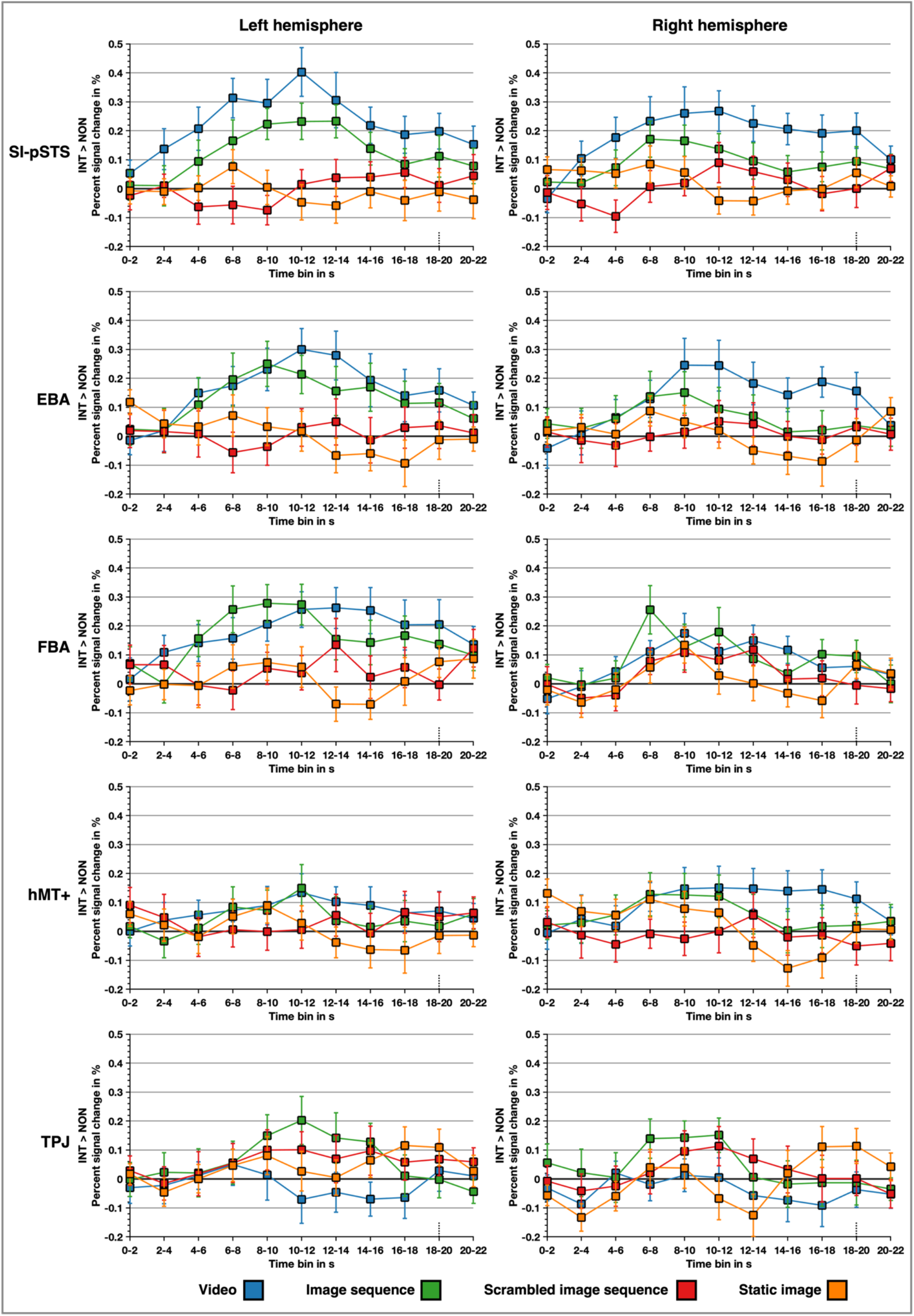
Illustration of bilateral ROI time course data displaying the PSC difference score (response to interactive minus non-interactive scenes) for each motion condition. Stimulus presentation occurred during the first nine time bins and responses blocks spanned the last two time bins.

**Table 1.** Behavioural rating means and standard deviations for interactive and non-interactive static stimuli

| Scene | How social? | How interesting? | How positive/negative? |
| --- | --- | --- | --- |
| *Interaction* | 4.54 (0.31) | 3.48 (0.45) | 0.49 (0.15) |
| *Non-Interaction* | 2.22 (0.47) | 2.68 (0.36) | -0.14 (0.12) |
| Mean difference | 2.32 | 0.80 | 0.63 |

**Table 2.**
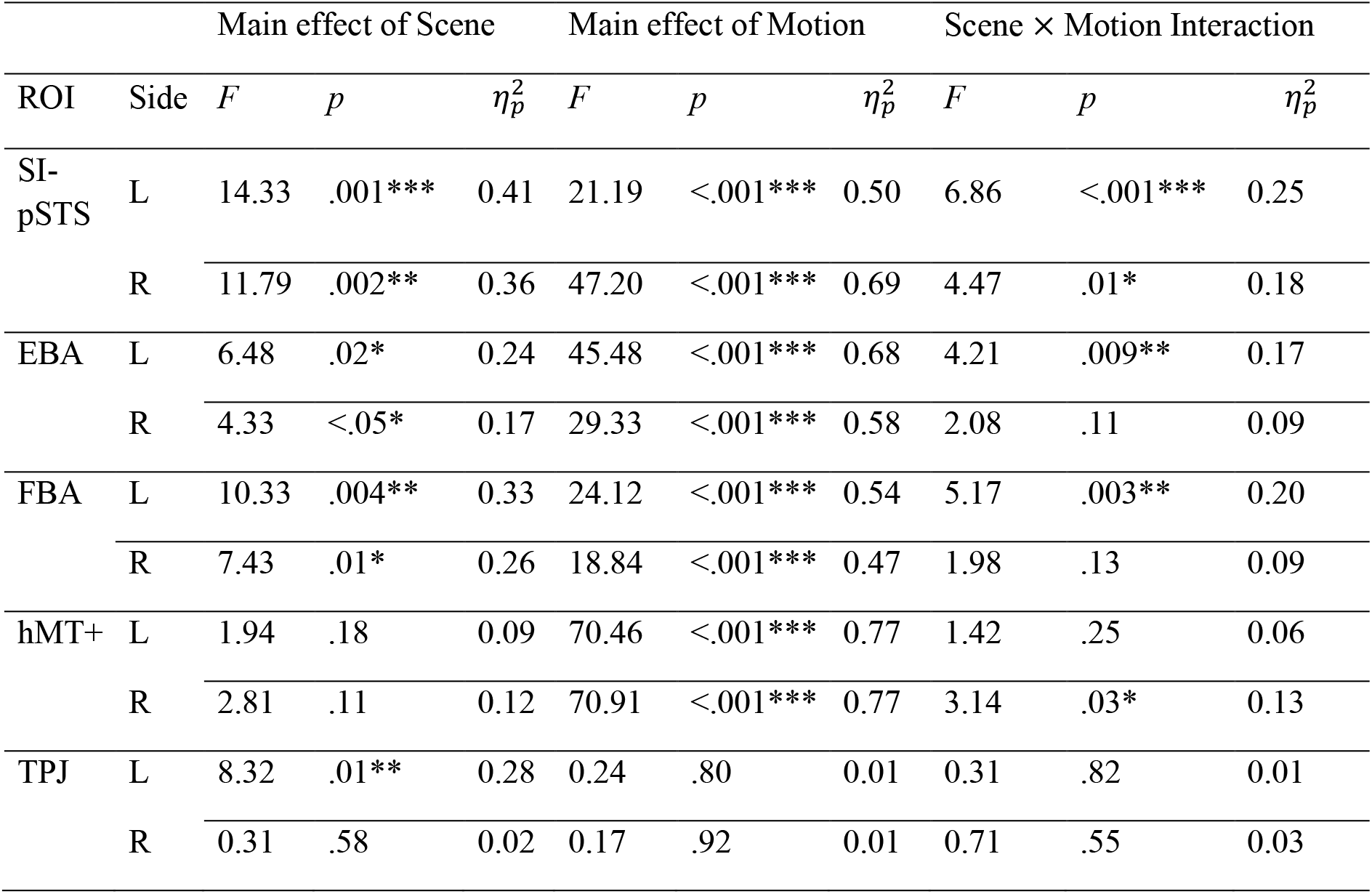
Inferential statistics of the 2 (Scene) × 4 (Motion) ANOVA by region of interest and hemisphere

| ROI | Side | Main effect of Scene | | | Main effect of Motion | | | Scene × Motion Interaction | | |
| --- | --- | --- | --- | --- | --- | --- | --- | --- | --- | --- |
| | | $F$ | $p$ | $\eta_p^2$ | $F$ | $p$ | $\eta_p^2$ | $F$ | $p$ | $\eta_p^2$ |
| SI-pSTS | L | 14.33 | .001*** | 0.41 | 21.19 | <.001*** | 0.50 | 6.86 | <.001*** | 0.25 |
|  | R | 11.79 | .002** | 0.36 | 47.20 | <.001*** | 0.69 | 4.47 | .01* | 0.18 |
| EBA | L | 6.48 | .02* | 0.24 | 45.48 | <.001*** | 0.68 | 4.21 | .009** | 0.17 |
|  | R | 4.33 | <.05* | 0.17 | 29.33 | <.001*** | 0.58 | 2.08 | .11 | 0.09 |
| FBA | L | 10.33 | .004** | 0.33 | 24.12 | <.001*** | 0.54 | 5.17 | .003** | 0.20 |
|  | R | 7.43 | .01* | 0.26 | 18.84 | <.001*** | 0.47 | 1.98 | .13 | 0.09 |
| hMT+ | L | 1.94 | .18 | 0.09 | 70.46 | <.001*** | 0.77 | 1.42 | .25 | 0.06 |
|  | R | 2.81 | .11 | 0.12 | 70.91 | <.001*** | 0.77 | 3.14 | .03* | 0.13 |
| TPJ | L | 8.32 | .01** | 0.28 | 0.24 | .80 | 0.01 | 0.31 | .82 | 0.01 |
|  | R | 0.31 | .58 | 0.02 | 0.17 | .92 | 0.01 | 0.71 | .55 | 0.03 |

##### SI-pSTS

In line with our hypotheses, response in bilateral SI-pSTS was greater for interactive compared to non-interactive scenes and increased as a function of motion. These main effects were qualified by a significant interaction effect. Bilaterally, SI-pSTS showed significant sensitivity to interactive content only for video and image sequence conditions, as measured by higher response to the interactive than non-interactive scenes (image sequence: rSI-pSTS: *t*(21) = 2.74, *p* = .01, *d_rm_* = 0.24, lSI-pSTS: *t*(21) = 4.34, *p* < .001, *d_rm_* = 0.43), video: (rSI-pSTS: *t*(21) = 4.01, *p* < .001, *d_rm_* = 0.52, lSI-pSTS: *t*(21) = 4.48, *p* < .001, *d_rm_* = 0.58).

##### EBA

Again, in line with our hypotheses, response in bilateral EBA was sensitive to motion. Contrary to our original expectations, however, EBA also responded more strongly to interactive compared to non-interactive scenes. The interaction effect was only significant in left EBA, where PSC was greater for interactive vs. non-interactive scenes for image sequence (*t*(21) = 2.91, *p* = .008, *d_rm_* = 0.28) and video (*t*(21) = 3.32, *p* = .003, *d_rm_* = 0.25) conditions but not static images or scrambled image sequence conditions (all *t*s < 0.63, all *p*s ≥ .53, *d_rm_* < 0.08). Nonetheless, planned contrasts in right EBA revealed significantly greater responses to interactive than non-interactive videos (*t*(21) = 3.44, *p* = .002, *d_rm_* = 0.18), but this difference was not significant for static images (*t*(21) = 0.89, *p* = .38, *d_rm_* = 0.09).

##### FBA

Bilateral FBA response was greater for dynamic stimuli *and* higher to interactive than non-interactive scenes. The interaction effect was only significant in left FBA, however, where PSC was greater for interactive vs. non-interactive image sequences (*t*(21) = 4.32, *p* < .001, *d_rm_* = 0.59) and video (*t*(21) = 3.80, *p* < .001, *d_rm_* = 0.53) conditions but not static images or scrambled image sequence conditions (all *t*s < 0.91, all *p*s ≥ .37, *d_rm_* < 0.14).

##### hMT+

Bilateral hMT+ responded more strongly to dynamic than static stimuli. For right hMT+, there was also an unexpected significant interaction effect; PSC was greater for the interactive vs. non-interactive scene in the video condition only (*t*(21) = 3.17, *p* = .005, *d_rm_* = 0.19; all other conditions *t*s < 1.18, all *p*s ≥ .25, *d_rm_* < 0.10).

##### TPJ

No significant effects were found for rTPJ, whereas lTPJ only showed a greater response to interactive compared to non-interactive scenes across motion conditions.

#### 3.2.2. Follow-up interaction selectivity analyses

Analyses of significant interactions between scene type and stimulus type (see Table 1) followed a mixed approach of planned (videos vs. static images, and image sequences vs. scrambled sequences) and exploratory (videos vs image sequences) comparisons. Specifically, we explicitly compared the differences in size of interaction specific responses between different motion conditions. A region sensitive to dynamic interactions should not only show greater interaction selectivity for videos than static images but also for videos compared to (implied motion) image sequences. Furthermore, an interaction specific region should be sensitive to the differences between meaningful vs scrambled image sequences. Originally, planned analyses only focused on bilateral SI-pSTS and EBA, however, PSC analyses revealed an interaction in left FBA, which was therefore included in the analyses here. Right hMT+ was not included in this follow-up analysis as it only showed a small effect of interaction selectivity for videos in the absence of an overall main effect of scene.

##### SI-pSTS

Bilaterally, interaction-selectivity was significantly greater for videos than static images (rSI-pSTS: *t*(21) = 3.51, *p* = .002, *d_rm_* = 0.86, lSI-pSTS: *t*(21) = 4.30, *p* < .001, *d_rm_* = 0.94). Interaction-selectivity was greater for coherent sequences of static image sequences than for the scrambled sequences only in the left SI-pSTS (*t*(21) = 2.16, *p* = .04, *d_rm_* = 0.56), and only rSI-pSTS showed a trend for greater interaction-selectivity for videos compared to image sequences (*t*(21) = 2.01, *p* = .06, *d_rm_* = 0.59). Qualitatively, these findings also match the pattern of time courses seen in bilateral SI-pSTS.

##### EBA

Interaction-selectivity in bilateral EBA was significantly greater for videos than static images (rEBA: *t*(21) = 2.71, *p* = .01, *d_rm_* = 0.65, lEBA: *t*(21) = 3.75, *p* = .001, *d_rm_* = 0.96). In contrast, bilaterally, EBA was equally interaction-selective for image sequences and scrambled image sequences (rEBA: *t*(21) = 0.43, *p* = .67, *d_rm_* = 0.12, lEBA: *t*(21) = 1.15, *p* = .26, *d_rm_* = 0.38), and for videos and image sequences (rEBA: *t*(21) = 1.49, *p* = .15, *d_rm_* = 0.40, lEBA: *t*(21) = 0.54, *p* = .59, *d_rm_* = 0.17). Time courses for EBA show a weaker, but similar pattern to SI-pSTS – particularly on the left.

##### FBA

Interaction-selectivity in left FBA was greater for videos than static images (*t*(21) = 2.78, *p* = .01, *d_rm_* = 0.67), as well as for image sequence than scrambled image sequences (*t*(21) = 2.53, *p* = .02, *d_rm_* = 0.62). Selectivity for videos was *comparable* to image sequences (*t*(21) = −0.30, *p* = .77, *d_rm_* = −0.07). Time courses match these results with selectivity primarily for meaningful image sequences and to some extent also for videos.

#### 3.2.3. Multivariate analyses

Here, multiple regression was used in an exploratory, data-driven approach to model interaction-selectivity as a voxelwise function of body-, motion-, and point-light-interaction selectivity as measured in the localisers (see Figure 3 Figure 4). This analysis examined whether each of these variables was a significant independent predictor of interaction selectivity in the main experiment. We focused in particular on the interactivity in the image sequence and video conditions because these revealed significant univariate effects of interaction vs non-interaction conditions. As described above, these analyses were performed participant-wise over a global bilateral ROI (union of SI-pSTS-EBA-hMT+; see Figure 3). The resulting regression betas were aggregated across subjects and tested against zero (see Figure 7 and Appendix 5, Table A5 for statistics).

**Figure 7.**
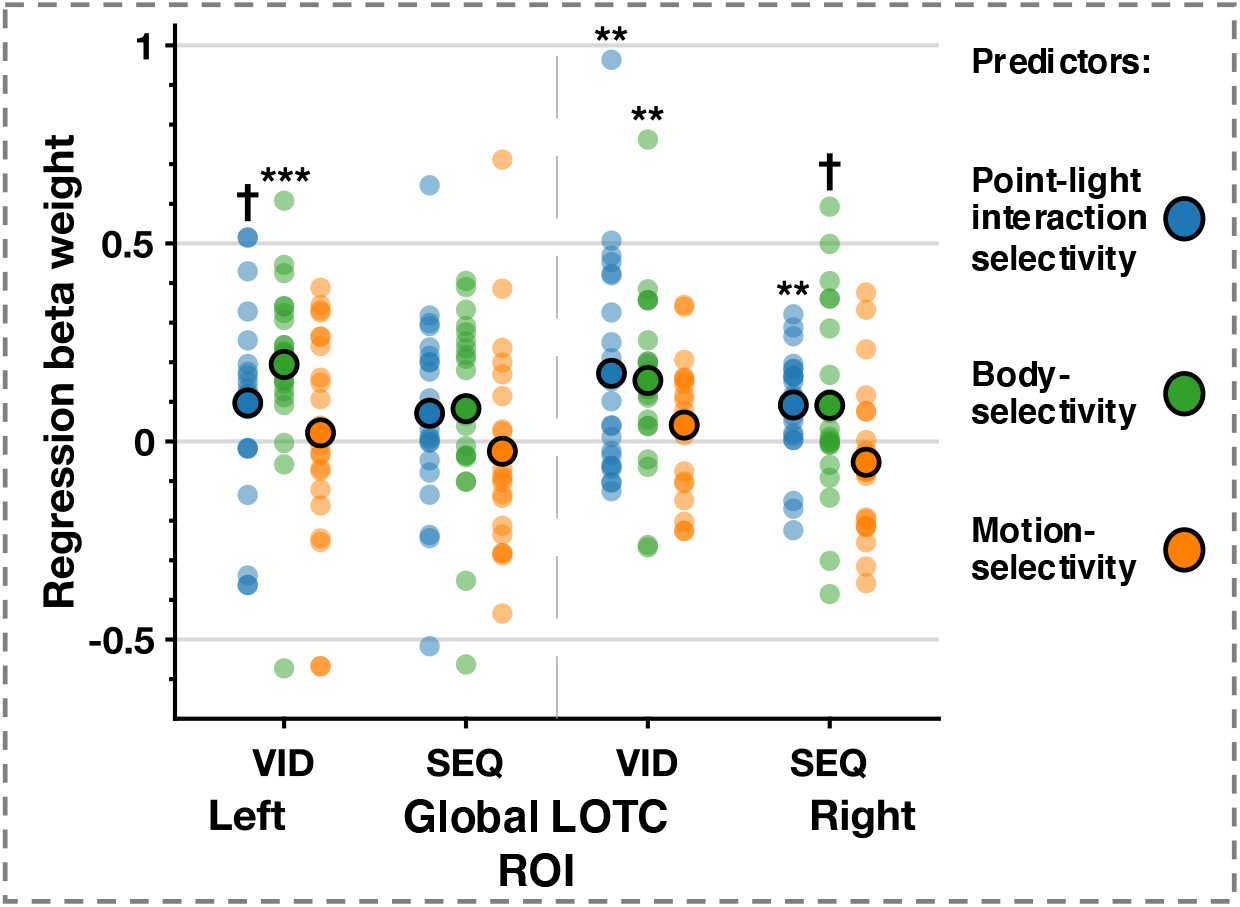
Sample mean and distribution of regression beta weights for each predictor (from left to right point-light-interaction-selectivity, body-selectivity, and motion-selectivity) predicting interaction-selectivity of videos (VID) and image sequences (SEQ) in the global ROI by hemisphere. Significant results of one sample t-test against zero are marked with *** (p < .0001), ** (p < .01), and † (marginal p < .10)

Interaction-selectivity in **videos** was significantly predicted bilaterally by body- and point-light interaction-selectivity (albeit only marginally in the left hemisphere), but not motion-selectivity. Interaction-selectivity in **image sequences** could be marginally predicted by body-, and point-light-interaction-selectivity, but not motion-selectivity. In simpler terms, voxels that tended to be more strongly body-selective, or more strongly selective to interactions expressed in point-light walkers, were more likely also to be selective for social interactions in the main experiment.

Importantly, although the univariate analyses revealed a stronger response to full-motion videos than to image sequences in nearly every ROI, when examined here at the level of voxel-wise patterns, low-level motion selectivity *per se* appears to contribute very little to the distributed response to social interactions. In simpler terms, strongly motion-selective voxels are no more likely to be interaction-selective than weakly-selective voxels. This finding goes some way to alleviate concerns that motion confounds between interacting and non-interacting videos may drive univariate differences in the response to social interactions, because we would expect such confounds to be even more apparent in multivoxel as opposed to univariate analyses.

## 4. Discussion

The present study investigated the contribution of motion sensitivity to social interaction perception within social brain regions involved in processing complex social scenes. Combining univariate and multivariate analyses, we confirm that bilateral SI-pSTS plays a central role in *dynamic* social interaction perception, but much less so when ‘interactiveness’ is conveyed solely with static cues. In addition, we find that EBA shows a similar profile. While SI-pSTS is somewhat more tuned to video interactions than is EBA, both bilateral SI-pSTS and EBA showed a greater response to social interactions compared to non-interactions and both regions responded more strongly to videos than static images. Crucially, interaction-selectivity in bilateral SI-pSTS and left EBA was motion-dependent, providing partial support for our key hypothesis. Specifically, these regions showed increased activation in response to social interactions compared to non-interactions when shown as videos and coherent image sequences – but not when motion cues were absent (in scrambled and static conditions). Exploratory multivariate regression analyses suggest that selectivity for simple visual motion does not in itself drive interactive sensitivity in either SI-pSTS or EBA. Rather, selectivity for interaction dynamics (captured here in the response to point-light animations), and selectivity for static images of bodies, make positive and independent contributions to this effect across the LOTC region.

Overall, our results are in line with prior work demonstrating the sensitivity of the SI-pSTS to dynamic social interactions, including prior work using static social scenes that implied motion (Kujala et al., 2012) and work comparing realistic vs rigid social movement (Georgescu et al., 2014). Further, our results are also congruent with Hafri et al. (2017) finding that the SI-pSTS showed no difference in activation to static scenes of social interaction versus non-interactions, although it should be noted that Hafri et al. used different scenarios in dynamic versus static conditions, unlike the current study. The sensitivity of the SI-pSTS to both interaction and motion is in stark contrast to the response in nearby TPJ, which is neither driven by these rich social stimuli, nor sensitive to manipulations of either interactivity or motion. While some prior work has suggested that TPJ is involved in the processing of complex scenes of social interaction (e.g., Canessa et al, 2012), it seems likely that this occurs primarily when social scenes or the task require mentalising (Masson & Isik, 2021; Walbrin et al., 2018), rather than when such scenes are unambiguous and participants are performing a non-social orthogonal task, as in the current work.

Similarly, our results in EBA are in line with some previous reports that EBA is sensitive to dynamic interactive information – at least when stimuli depict human bodies (Abassi & Papeo, 2020; Bellot et al, 2021; Walbrin & Koldewyn, 2019; though see Walbrin et al, 2020). In contrast to previous work (Abassi & Papeo, 2020, 2021), however, in our results EBA did not exhibit preferential activation for social interactions versus non-interaction when scenes were static. Crucially, the stimuli used in the current work were naturalistic, every-day social scenes that included a variety of interactive cues. In contrast, the focus of Abassi & Papeo’s work has been on ‘prototypical’ social interactions and a single interactive cue – that of facing direction. Indeed, their proposal is that the visual system is tuned for quick and accurate perception of face-to-face (vs. non-facing) bodies (Papeo, 2020), something they suggest could take place primarily in EBA. If true, the previously reported interaction selectivity in EBA to static stimuli could primarily rely on facing direction, explaining why such selectivity is not seen in our results, which rely on a richer set of interactive cues.

Our findings in SI-pSTS and EBA are in line with recent proposals that there is a third visual stream, which is specialised for conveying dynamic social information from V1, through hMT+ and EBA to the STS (Pitcher, 2021; Pitcher & Ungerleider, 2021). This picture is somewhat complicated, however, by our unexpected result that FBA, a region firmly in the ventral stream (Schwarzlose et al, 2008), is also more responsive to the video condition *and* shows some sensitivity to interactive information, particularly in the left hemisphere. However, given widespread connectivity between visual streams and amongst person-selective regions (see Donato et al, 2020 for a review), FBA’s response profile is perhaps not so surprising. As has been shown previously for biological motion (Dasgupta et al, 2017; Duarte et al, 2022), it seems likely that ventral stream regions would contribute form information to the pSTS during dynamic social interaction processing, and that response in ventral regions would increase with the complexity of social scenes (Haxby et al, 2020) . In line with our original hypotheses, however, we would still expect ventral regions to be less sensitive to our key experimental manipulations. Indeed, while FBA statistically shows sensitivity to both motion and interactive information, a look at the time-course responses (Figure 6) casts some doubt about the strength and consistency of these differences. Left FBA shows an earlier, more consistent, and more interaction sensitive response to the image *sequence* than to the video condition, while there is little evidence of consistent differences in interaction sensitivity across conditions in right FBA. This is consistent with prior work showing that while FBA does respond to dynamic bodies, including point-light displays (Atkinson et al., 2012; Peelen et al., 2006), it shows considerably less variability in response between dynamic and static displays than either the EBA or face selective pSTS (Pitcher et al., 2019). Most prior work on interaction perception has not focused on, or indeed even considered, the FBA, and whole-brain results across studies for dynamic interactions suggest that FBA does not typically show strong interaction selectivity (Isik et al, 2017; Walbrin et al, 2018, Walbrin et al, 2020), especially in the absence of full body stimuli. There is, however, some recent work that reports greater activation in FBA in response to static pictures of facing vs non-facing dyads (Abassi & Papeo, 2021). In light of these mixed findings, it is difficult to draw any strong conclusions regarding FBA’s sensitivity to (especially dynamic) interactive information from our data. However, our results suggest that future work needs to take a careful look at FBA’s role in social interaction and social scene processing as well as thinking more globally about network contributions to social interaction processing.

One strength of the current work is the use of naturalistic, yet controlled stimuli. The dynamic and static conditions used the same scenarios, and interactive vs. non-interactive conditions were well-matched for motion energy, the action depicted, scenic and object elements, and actor identity. As an even tighter control, intact image sequences where scenes show meaningful change over time used the same set of pictures as the scrambled sequences, where implied motion cues (and meaning) are disrupted. Similarly, the richness and “naturalness” of the stimuli are also a strength, allowing us to probe social interaction perception as it might occur in the ‘real world’, where social interactions take place within a wider context and rely on a mix of visual cues. The richness of our stimulus set also creates interpretation challenges, however. As in real life, participants found observed social interactions to be both more interesting and more positive than corresponding non-interactions. While the mean differences between interactions and non-interactions for both visual interest and valence were small, they were reliable. Thus, it is possible that some of the difference in brain response to interactions and non-interactions could be at least partially driven by heightened interest and/or increased attention to human information in scenes of social interactions (Skripkauskaite et al., 2021). Similarly, despite the use of a task that did not require mentalising or social processing, our results could reflect top-down influence from the mentalising network, as participants are intuitively more likely to engage in mentalising when viewing social interactions. We think this unlikely, however, given that bilateral TPJ did not show sensitivity to the difference between interactive and non-interactive scenes in *any* motion condition. In addition, our whole brain results (Appendix 6) do not suggest wide-spread activity differences between interactive and non-interactive conditions (collapsed across motion conditions) in either the attention or mentalising networks. As a result, we think it unlikely that the relatively small differences in perceived interest or valence are driving the whole of our results. This is particularly true considering the fact that our results are congruent with prior work using less naturalistic stimuli. That humans find naturalistic interactions particularly engaging is a feature of our intrinsically social nature, and it may be nigh impossible to capture natural social interaction perception in the absence of heightened social interest.

## 5. Conclusions

Using naturalistic social scenes and manipulating both interactive content and motion information, we found that the SI-pSTS, a region that may be uniquely engaged by “social interactiveness” across diverse cues and stimulus types, is highly reliant on *dynamic* interactive information. In line with the proposal that dynamic information for social understanding is preferentially processed in the ‘third visual stream’ (Pitcher & Ungerleider, 2020), EBA shows a similar – if less selective – response profile. Our results strongly suggest that – at least when interactive information is conveyed primarily via *body* information – EBA and SI-pSTS work together, in line with recent work showing that information flow between EBA and SI-pSTS increases when facing dyads move towards rather than away from each other (Bellot et al, 2021). Our results are also in line with proposals that the pSTS is the “hub” of the dynamic social perception system, that flexibly integrates relevant information from earlier social perception regions during interactive processing. Indeed, our results suggest that two independent processes, that are not strictly segregated within focal regions, may underlie similar response profiles across SI-pSTS and EBA. One process may be picking up on dynamic social cues (reflected in selective responses to point-light interactions), while the other is more focused on static form cues (reflected in selective responses to static body postures). Our findings motivate future work looking at the integration of social information across visual streams, and at how, and when our social perception systems become tuned to social interactions across development.

## Data and Code Availability Statement

In-house code and stimulus examples as well as the data that support the findings of this study are openly available at the project’s OSF page, at this <u>link</u>.

## Author contributions

**Julia Landsiedel**: Methodology, Investigation, Formal analysis, Project administration, Visualization, Writing – Original Draft, Writing – Review & Editing; **Katie Daughters**: Conceptualization, Methodology, Investigation, Formal analysis, Project administration, Writing – Original Draft, Writing – Review & Editing; **Paul Downing**: Methodology, Writing – Original Draft, Writing – Review & Editing; **Kami Koldewyn**: Conceptualization, Funding acquisition, Methodology, Investigation, Supervision, Writing – Original Draft, Writing – Review & Editing

## Acknowledgements

We are grateful to Laura Jastrzab for her help with data collection as well to members of the Social Neuroscience and Cognition group and Bangor Imaging Group at Bangor University for general feedback, helpful discussion, and suggestions throughout the research process.

