## Supplementary Information for "The role of motion in the neural representation of social interactions in the posterior temporal cortex"

**Appendix 1: Stimulus set validation study**

To validate the stimulus set used in this study, 55 participants (mean age = 29.87, SD = 9.03, 18 male) participated in an online study also created using the PsychoPy (Peirce et al., 2019) and hosted via Prolific (https://www.prolific.co). Participants were shown all videos and the static image key frame from each video. They were asked to rate each stimulus on three factors: 1) to what extent each image/video depicted a social interaction (on a 5-point Likert scale from "definitely not" to "definitely"); 2) how visually interesting was the image/video (on a 5-point Likert scale from "very un-interesting" to "very interesting"; and 3) how positive/negative they thought the image/video was (on a 3-point Likert scale from "negative" to "positive").

Three paired *t*-tests were carried out on the video- and image-ratings respectively (for descriptive statistics please see Table A1). Results confirmed that 1) social interactions were rated as more socially interactive compared to the non-interaction stimuli (Videos: $t(54) = 20.039$, $p < .001$; Images: $t(54) = 19.836$, $p < .001$); 2) social interactions were rated as more visually interesting compared to non-interaction stimuli (Videos: $t(54) = 8.918$, $p < .001$; Images: $t(54) = 9.971$, $p < .001$); and 3) that social interactions were rated as more positive compared to non-interactions (Videos: $t(54) = 15.650$, $p < .001$; Images: $t(54) = 12.685$, $p < .001$).

*Table A1. Rating means (SD) for interactive and non-interactive videos and static image stimuli from the validation study.*

| Motion condition | Scene | How social? | How interesting? | How positive/negative? |
|---|---|---|---|---|
| Video | Interaction | 4.39 (0.54) | 3.07 (0.81) | 0.52 (0.24) |
| | Non-Interaction | 1.94 (0.65) | 2.28 (0.67) | -0.16 (0.15) |
| | *Mean difference* | *2.45* | *0.79* | *0.68* |
| Image | Interaction | 4.25 (0.46) | 2.82 (0.69) | 0.35 (0.18) |
| | Non-Interaction | 2.07 (0.67) | 2.13 (0.64) | -0.06 (0.14) |
| | *Mean difference* | *2.18* | *0.69* | *0.41* |

**Appendix 2: Motion energy calculations**

To assess potential differences in motion energy for the videos of the interactive and non-interactive scenes, we followed the approach by Grezes et al. (2007). Motion energy for each video was calculated based on light intensity variation and averaged across pixels that scored higher than a noise level of 10 using a custom Matlab script, which can be accessed here.

**Appendix 3: ROI definition**

*Table A2. MNI coordinates (based on group level localizer task contrasts) used as centre of 8mm bounding spheres in initial step to subject-specifc ROIs defined at an uncorrected threshold of p < .01*

| Region | Task | Defining contrast | Right | Left |
|---|---|---|---|---|
| TPJ | Partly Cloudy localizer | Mentalizing > Pain | 48 -64 34 | -42 -68 36 |
| pSTS | Social interaction localizer | Interaction > Non-*i*nteraction | 56 -46 8 | -58 -50 8 |
| EBA | Body regions localizer | Bodies > Chair | 52 -60 -4 | -50 -68 6 |
| FBA | Body regions localizer | Bodies > Chair | 44 -42 -20 | -42 -48 -22 |
| hMT+ | Motion localizer | Moving rings > static rings | 46 -70 -4 | -46 -70 -2 |

### Appendix 4: Percent signal change descriptives for main and interaction effects

*Table A3. Overview of PSC 2 × 4 ANOVA estimated marginal means (SE) of main effects analyses*

| | | TPJ | | pSTS | | EBA | | hMT+ | | FBA | |
|---|---|---|---|---|---|---|---|---|---|---|---|
| | | **L** | **R** | **L** | **R** | **L** | **R** | **L** | **R** | **L** | **R** |
| **Scene** | *Interaction* | 0.30 (0.09) | 0.20 (0.06) | 0.53 (0.10) | 0.74 (0.10) | 0.94 (0.14) | 0.95 (0.15) | 0.89 (0.11) | 1.21 (0.11) | 1.01 (0.07) | 0.95 (0.13) |
| | *Non-Interaction* | 0.22 (0.08) | 0.18 (0.06) | 0.37 (0.09) | 0.60 (0.09) | 0.83 (0.13) | 0.86 (0.14) | 0.83 (0.11) | 1.16 (0.11) | 0.87 (0.08) | 0.84 (0.11) |
| **Motion** | *Static image* | 0.27 (0.10) | 0.18 (0.07) | 0.24 (0.06) | 0.29 (0.08) | 0.40 (0.10) | 0.42 (0.09) | 0.31 (0.08) | 0.55 (0.06) | 0.74 (0.07) | 0.69 (0.10) |
| | *Scrambled image sequence* | 0.25 (0.09) | 0.20 (0.06) | 0.42 (0.10) | 0.69 (0.10) | 0.86 (0.13) | 0.91 (0.16) | 0.82 (0.11) | 1.13 (0.11) | 0.99 (0.08) | 0.93 (0.12) |
| | *Image sequence* | 0.28 (0.07) | 0.20 (0.06) | 0.45 (0.10) | 0.68 (0.11) | 0.85 (0.14) | 0.90 (0.14) | 0.83 (0.11) | 1.18 (0.11) | 0.98 (0.08) | 0.94 (0.13) |
| | *Video* | 0.24 (0.09) | 0.18 (0.06) | 0.69 (0.11) | 1.01 (0.11) | 1.43 (0.19) | 1.37 (0.22) | 1.49 (0.15) | 1.89 (0.18) | 1.07 (0.08) | 1.04 (0.13) |

*Table A4. PSC Condition means (SE) for the main experimental task*

| | | TPJ | | pSTS | | EBA | | hMT+ | | FBA | |
|---|---|---|---|---|---|---|---|---|---|---|---|
| **Scene** | **Motion** | **L** | **R** | **L** | **R** | **L** | **R** | **L** | **R** | **L** | **R** |
| *Interaction* | *Static image* | 0.29 (0.11) | 0.19 (0.07) | 0.26 (0.07) | 0.31 (0.08) | 0.38 (0.10) | 0.44 (0.09) | 0.30 (0.08) | 0.52 (0.06) | 0.77 (0.07) | 0.72 (0.11) |
| | *Scrambled image sequence* | 0.28 (0.11) | 0.20 (0.07) | 0.45 (0.11) | 0.75 (0.10) | 0.88 (0.15) | 0.93 (0.18) | 0.85 (0.13) | 1.15 (0.12) | 1.02 (0.08) | 0.95 (0.14) |
| | *Image sequence* | 0.34 (0.10) | 0.25 (0.07) | 0.56 (0.11) | 0.74 (0.11) | 0.94 (0.14) | 0.94 (0.14) | 0.86 (0.11) | 1.21 (0.10) | 1.09 (0.07) | 1.03 (0.14) |
| | *Video* | 0.27 (0.09) | 0.16 (0.07) | 0.86 (0.13) | 1.15 (0.12) | 1.55 (0.20) | 1.47 (0.23) | 1.55 (0.15) | 1.97 (0.18) | 1.17 (0.08) | 1.10 (0.14) |
| *Non-Interaction* | *Static image* | 0.24 (0.09) | 0.17 (0.07) | 0.22 (0.07) | 0.27 (0.09) | 0.43 (0.11) | 0.40 (0.09) | 0.32 (0.09) | 0.57 (0.07) | 0.72 (0.08) | 0.65 (0.10) |
| | *Scrambled Image sequence* | 0.22 (0.10) | 0.19 (0.07) | 0.39 (0.11) | 0.64 (0.11) | 0.83 (0.13) | 0.89 (0.14) | 0.80 (0.11) | 1.11 (0.10) | 0.96 (0.09) | 0.91 (0.11) |
| | *Image sequence* | 0.22 (0.06) | 0.16 (0.06) | 0.34 (0.09) | 0.62 (0.11) | 0.76 (0.14) | 0.86 (0.15) | 0.79 (0.12) | 1.15 (0.12) | 0.86 (0.09) | 0.84 (0.11) |
| | *Video* | 0.22 (0.09) | 0.21 (0.05) | 0.53 (0.11) | 0.86 (0.11) | 1.30 (0.18) | 1.27 (0.21) | 1.42 (0.15) | 1.82 (0.18) | 0.96 (0.09) | 0.98 (0.12) |

**Appendix 5: Multivariate regression analysis statistics**

*Table A5. Results of the one-sample t-tests against zero on the mean beta weights for each of three predictors of the multiple regression analyses on motion-dependent interaction-selectivity in the pSTS-EBA-hMT+ union ROI by hemisphere. Significant p-values are in bold, marginally significant p-values in italic.*

| | | *Predictors* | | | | | | | | |
|---|---|---|---|---|---|---|---|---|---|---|
| | | **Point-light-interaction selectivity** | | | **Body-selectivity** | | | **Motion-selectivity** | | |
| **Hemisphere** | **Interaction selectivity condition** | *beta* | *t*(21) | *p* | *beta* | *t*(21) | *p* | *beta* | *t*(21) | *p* |
| Left | Video | 0.10 | 1.85 | *<.10* | 0.19 | 4.01 | **<.001** | 0.02 | 0.38 | 0.71 |
| | Image sequence | 0.07 | 1.38 | 0.18 | 0.08 | 1.65 | 0.11 | -0.03 | -0.45 | 0.65 |
| Right | Video | 0.17 | 2.91 | **<.01** | 0.15 | 3.24 | **<.01** | 0.04 | 1.18 | 0.25 |
| | Image sequence | 0.09 | 3.04 | **<.01** | 0.09 | 1.75 | *<.10* | -0.05 | -1.26 | 0.22 |

**Appendix 6: Whole-brain analyses**

Behavioural data indicated that interactive vs non-interactive scenes significantly differed for both visual interest and valence (see main text behavioural results), suggesting potential attentional differences between conditions. In addition, such differences could lead to differential activity within the mentalising networks. To examine these differences on a neural level, the first-level t-maps for the contrasts of interest (interactive vs non-interaction scene across motion conditions respectively) were entered into a second level random effects model using one-sample t-test for group analyses. The resulting group level t-map were height-thresholded at $p < .001$ and false discovery rate cluster corrected at $p < .05$ (cluster extent threshold k = 177).

The two surviving clusters (see also Figure S1) were located in (1) a region encompassing the right putamen and caudate nucleus (MNI peak coordinate x = 26, y = 16, z = -6, peak t-value = 4.88, cluster size = 377), as well as (2) a region encompassing the left posterior superior temporal sulcus and middle temporal gyrus (MNI peak coordinate x = -62, y = -52, z = 14, peak t-value = 4.74, cluster size = 177).

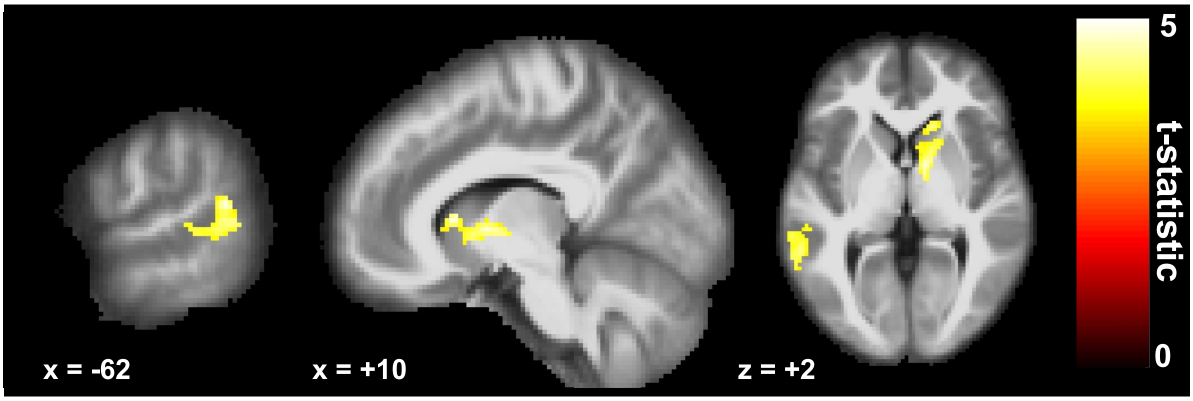


*Figure A1. Sagital and axial slices illustrating cluster locations of the whole-brain group level contrast of interactive vs non-interactive scenes p < .001 k = 177, equivalent of $p_{FDR} < .05$.*
